# Engineering drive-selection balance for localised population suppression with neutral dynamics

**DOI:** 10.1101/2024.05.21.595228

**Authors:** Katie Willis, Austin Burt

## Abstract

Whilst the release of sterile males has been highly successful in suppressing some pest populations, it is impractical for other species due to the males disappearing after a single generation, necessitating large, repeated releases to maintain sufficient impact. Synthetic gene drives promise more efficient approaches since they can increase in frequency from rare, yet this also allows them to spread across a landscape, which may not always be desired. Between these two extremes are selectively neutral genetic constructs which persist at the frequency they are released, offering the potential for efficient suppression that remains localised. One way to achieve this would be to have perfect balance, at all construct frequencies, between gene drive increasing frequency and selection decreasing it. Here we describe a way to create this balance involving a toxin-antidote genetic construct that causes recessive lethality, encodes a genomic editor that makes dominant lethal edits in the genome, and provides protection against the action or consequences of the editing. Computer modelling shows that this design can be 100-fold more efficient than sterile males, increasing to 1000-fold more when released alongside a genetic booster. We describe designs for CRISPR-based molecular construction, including options that avoid using recoded genes as antidotes.

## 1. Introduction

Pest populations continue to impose a substantial health and economic burden on humanity by transmitting vector borne diseases, harming crops, and causing unwanted environmental change. Recent advancements in genome engineering technologies have facilitated the development of a variety of genetic approaches for controlling pest populations. These involve releasing modified members of the pest species designed to mate with those in the wild and reduce harm (Grilli et al., 2021, Raban et al., 2023). One approach is for the released individuals to suppress the density of the population by interfering with its ability to successfully reproduce, thereby reducing the overall harm caused. Of these, the most widely used strategy is the sterile insect technique (SIT), where the released males are sterilised through irradiation (Dyck et al., 2021), or related techniques where males are effectively sterilised by *Wolbachia* cytoplasmic incompatibility (Mohanty et al., 2016, Ching, 2021, Crawford et al., 2020) or more precise genetic modifications (Alphey, 2002, Kandul et al., 2019). This approach has been successful in supressing some populations (Hendrichs et al., 2002, Vreysen et al., 2000, Wyss, 2000), but significant and sustained suppression requires repeat, inundative releases, making it impractical for large target populations or for species difficult to rear.

One reason for the inefficiency of sterile male releases is that the causal agent does not persist from generation to generation, and thus repeated releases are required to maintain impact. The application of selfish genetic elements that spread from low frequency (genetic drive) has been proposed as one way to increase efficiency by reducing the numbers one needs to release (Burt, 2014, Grilli et al., 2021, Hay et al., 2021). Here, smaller releases are required because the genetic construct not only can persist from generation to generation, but moreover can increase in frequency and spread to neighbouring populations wherever there is gene flow. While this may be desirable in species that are harmful throughout their range, it may not always be appropriate if control is only desirable in part of the species range. In these latter cases it would be useful if the genetic element responsible for the reproductive load persisted over multiple generations, requiring smaller releases, but was unable to increase in frequency, preventing spread far from its release site. In the ideal case the genetic construct would persist at the frequency it was released at, being neither selected for or against, whilst nevertheless imposing a reproductive load upon the population – a selectively neutral suppressor (Burt and Deredec, 2018).

In this paper we propose and model one way to achieve this, by pairing a load-inducing construct with a toxin-antidote drive mechanism that perfectly counteracts the negative selection. We compare the efficiency of our design with alternate strategies for localisable suppression, assess its robustness to molecular imperfections and describe how it could be constructed using novel combinations of existing molecular tools for which there is ample precedent. Our modelling work presents the first step in the genetic biocontrol developmental pipeline and stimulates progression to the next stages: developing the approach in organisms of interest within the laboratory and evaluating their impact in the field.

## 2. Results

### 2.1. Engineering drive-selection balance

To reduce the number of individuals in a population one would need to introduce a genetic modification that reduces fitness. The most robust way to do this is to disrupt a gene which is needed for survival or reproduction, and disruption of most essential genes results in recessive fitness effects. Alleles with reduced fitness would usually decline in frequency due to natural selection, and for a recessive allele selection would be proportion to its frequency, being very small when rare, since the allele is most often in heterozygotes where there are no fitness costs, and large when common, because the allele is more likely to be found in the homozygous state where it does not survive. One way to counteract this selection would be to pair a gene drive component with the recessive lethal allele that has a fitness advantage that perfectly balances the selection against it. A homing gene drive would be too strong, resulting in genetic drive at all frequencies and making the combination more suited as a low-threshold suppression strategy designed to spread across a landscape (Burt, 2003, Kyrou et al., 2018, Oberhofer et al., 2018, Hammond et al., 2021b, Yang et al., 2022). Rather, to perfectly balance a recessive lethal mutation, drive must be very weak at low frequency, increasing in strength in proportion to allele frequency. One class of gene drive mechanisms that can have this property are those based on toxin-antidote interactions (Hay et al., 2021, Ward et al., 2011, Wade and Beeman, 1994, Davis et al., 2001). The most promising and easiest to engineer are Cleave and Rescue systems that use CRISPR/Cas9 technology, in which Cas9 “cleaves” a target gene and creates an edited non-functional version which acts as a toxin and the antidote is a recoded version of the target gene that “rescues” function but is resistant to Cas9 cleavage (Champer et al., 2020b, Oberhofer et al., 2019).

To test whether a cleave-rescue drive would balance selection against a recessive allele we constructed an analytical population genetics model with three alleles: the wild-type, a genetic construct containing a genomic editor capable of editing the wild-type allele, and the edit created by the editor. In Fig. 1a we show that a cleave-rescue design involving a genomic editor that creates dominant lethal edits and is protected against the action or consequences of its own editing shows the desired dynamics (red line, where 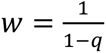) of precisely balancing the relative fitness of a fully recessive lethal mutation (orange line, where *w* = 1 − *q*) across the entire frequency range. As a result, when combined with recessive lethality the construct is selectively neutral (*w* = 1) across the full range of frequencies (Fig. 1a blue line). Despite the construct being selective neutrality, it still imposes a reproductive load (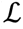) on the population that increases with frequency, where 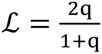 (Fig. 1b blue line). This load arises due to both the recessive lethality of the construct, acting to remove homozygotes (Fig. 1b orange line), and the action of the protected editor generating unprotected dominant lethal edits in heterozygotes (Fig. 1b red line).

**Fig. 1.**
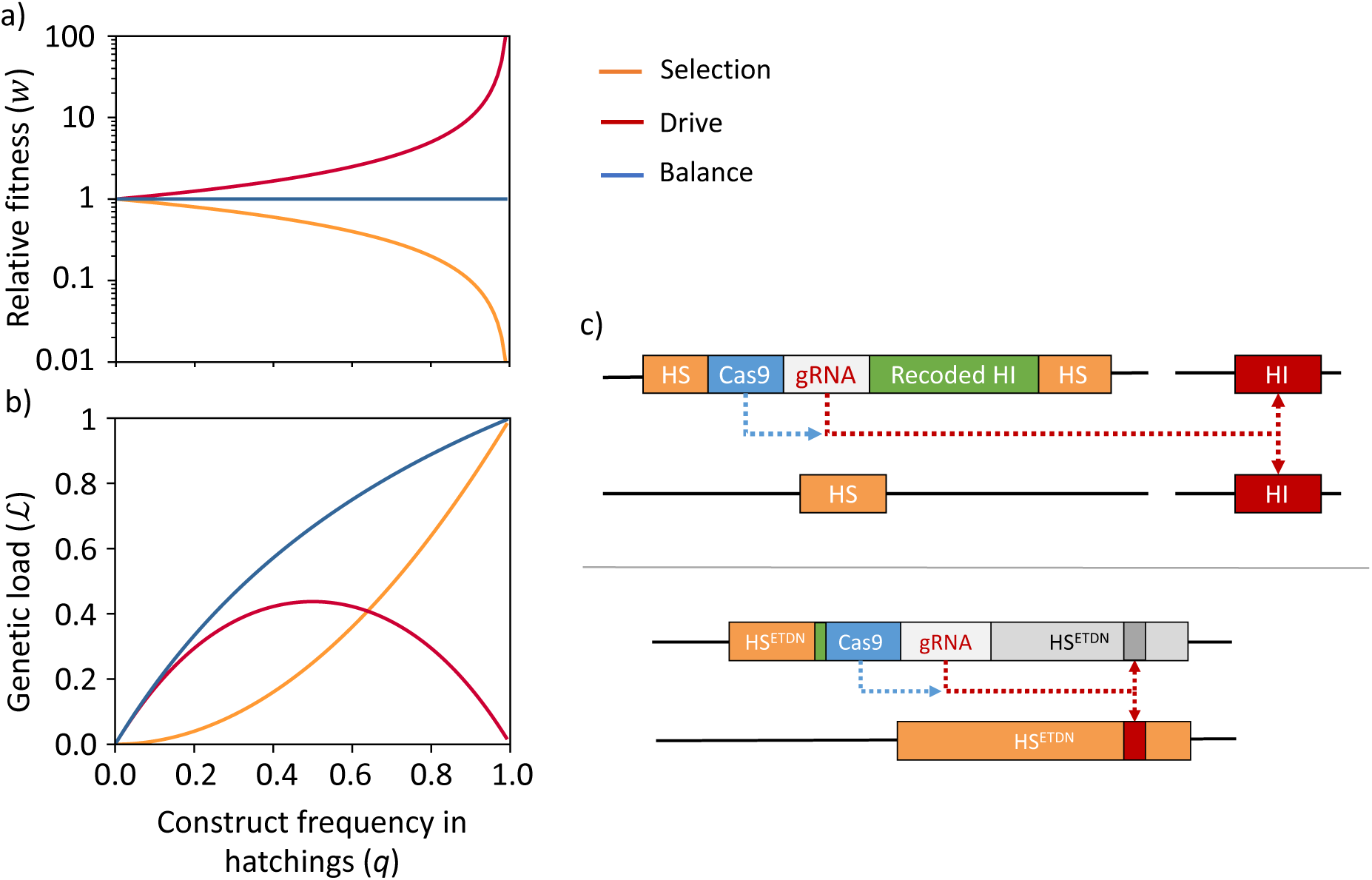
**(a)** The relative fitness (*w*) of different genetic constructs as a function of their frequency (*q*) in a population at Hardy-Weinberg equilibrium. The relative fitness of an allele which causes recessive lethality (orange) is comparable to wild type at low frequency and decreases as its frequency increases (*w* = 1 – *q*). In contrast, whilst a cleave-rescue based drive allele which creates dominant lethal mutations and provides protection against them also exhibits fitness comparable to the WT at very low frequency (red), its fitness increases with frequency due to drive 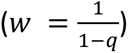. When combined (blue), the fitness advantage of the drive perfectly counteracts the fitness costs of the recessive lethal allele, resulting in fitness equal to the wild type at all frequencies (*w* = 1). Fitnesses are calculated from changes in construct frequency over a single generation, censused at the hatching stage: 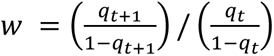 and *t* is the generation. **(b)** The genetic load (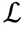) imposed on the population as a function of the frequency of the genetic construct. Load is calculated as the proportional reduction in reproductive output compared to a wild-type population over a single generation. **(c)** Possible CRISPR/Cas9 molecular configurations. Recessive lethality can be achieved by inserting the construct into a haplo-sufficient gene (HS, orange box) required for survival in both sexes such that it disrupts function. The upper panel illustrates how the cleave-rescue drive can be implemented using two genes, in which the genetic construct encodes a Cas9 and gRNA which generates non-functional edits in a haplo-insufficient gene (HI, red box) during gametogenesis. Protection can be achieved by incorporating a recoded version of the HI gene into the construct that cannot be edited by the gRNA but is still able to fully function (Recoded HI, green box). The lower panel shows an alternative implementation involving a single HS gene that contains one or more sites that can be modified by the genomic editor to create dominant negative mutations (HS^ETDN^, orange box). If the construct is inserted upstream of the edit and generates a premature stop codon (green box) that prevents the dominant negative allele from being expressed the construct provides protection against edits in cis, without the need for a recoded copy of the gene. Note that although we have used CRISPR/Cas9 for illustration, alternative engineerable genome editors would also be suitable.

In principle, a genomic editor with the desired features could be built in multiple ways. Following from previous proposals (Oberhofer et al., 2019, Champer et al., 2020b), one could create a genetic construct that (1) is inserted into a haplo-sufficient (HS) gene required for survival in both sexes such that its presence disrupts function and creates recessive lethality, (2) encodes Cas9 and gRNA that act in the germline to create dominant knockout mutations in a haplo-insufficient (HI) gene required for survival and (3) contains a functional recoded version of the target gene resistant to editing that rescues the dominant fitness effects of a single copy of the edit (Fig. 1c upper). A simpler approach, not to our knowledge previously proposed, would be to insert the construct into a HS gene required for survival in both sexes which can also be edited by the genomic editor to create a dominant negative mutation. This kind of mutation creates a protein which interferes with the normal function of the wild-type protein, and therefore if expressed in heterozygotes would cause dominant lethality. In this design, if the construct is inserted such that its presence creates a premature stop codon upstream of the dominant negative edit, the construct will prevent expression of the edited allele located on the same chromosome, and thus provide protection against its dominant negative effects (Fig. 1c lower). This would obviate the need for the construct to contain a recoded version of the target gene, potentially making it significantly simpler to engineer. In this paper we explore the performance and robustness of this single-locus design, referred to as a protected dominant negative editor (PDNE), and further consider alternative two-locus designs in the supplementary material.

### 2.2. Performance

We next assess the performance of the PDNE by simulating releases of males heterozygous for the construct into a single, well-mixed population with two life stages, juveniles and adults, with density dependent mortality occurring at the juvenile stage. For comparison, a range of alternative self-limiting genetic strategies currently in development are also modelled, including the release of:

a) Males homozygous for dominant lethal genes in which:

I. the gene affects survival of both sexes before density dependent mortality, equivalent to the sterile male technique (SIT) (Oliva et al., 2012, Bellini et al., 2013) or individuals carrying *Wolbachia* leading to incompatibility (O’Connor et al., 2012, Mains et al., 2016).
II. the gene affects survival of both sexes acting after density dependent mortality, equivalent to a transgenic knockout of a gene required, for example, for pupal to adult maturation (RIDL) (Phuc et al., 2007, Harris et al., 2011, Lacroix et al., 2012, Carvalho et al., 2015).
III. the gene causes lethality in females only and acts after density dependent mortality (fsRIDL) (Fu et al., 2010, Labbé et al., 2012, Thomas et al., 2000).
b) Males homozygous for an autosomal X-shredder (XS) which causes sex-ratio distortion towards males (Galizi et al., 2014, Galizi et al., 2016, Pollegioni et al., 2020).
c) Males carrying a Y-linked editor (YLE) which creates dominant lethal female-specific edits, or an autosomal dominant lethal allele that causes lethality in females and drives via homing in males (fs-RIDL-drive) (Burt and Deredec, 2018).

For each of these strategies we modelled the idealised case of perfect genetic efficiencies and no unintended fitness effects. Figure. 2 shows the relative numbers of females in a population following repeated releases of modified males each generation at 10% or 50% of the original male population size. The combination of persistence and load of the PDNE means that it is substantially more efficient than SIT, RIDL, fsRIDL and XS at suppressing the population and is comparable to the previous best in class YLE or fs-RIDL-drive. At both release frequencies the PDNE can eliminate the population where the other strategies either cannot or require more generations to do so. This improvement in efficiency can be quantified by comparing the numbers of males needed to be released each generation to achieve a certain level of suppression. For example, releasing PDNE-carrying males each generation at 5.2% of the original male population size is sufficient to achieve 95% suppression within 36 generations in a population with an intrinsic rate of increase (*R_m_*) of 20, offering a 94-fold increase in efficiency compared to SIT. Improved efficiency is observed in populations with a range of different intrinsic growth rates and for a variety of suppression goals, where the level of suppression or timeframe in which the suppression is desired varies (Fig. 3, Table 1 and Supp. Fig. 1). The greatest reductions in release rate requirements compared to alternative strategies occur when there is a longer timeframe for the suppression goal to be achieved and in populations with higher *R_m_*. If the level of suppression required is lower, even greater gains in efficiency are observed (Table 1 and Supp. Fig. 1).

**Fig. 2.**
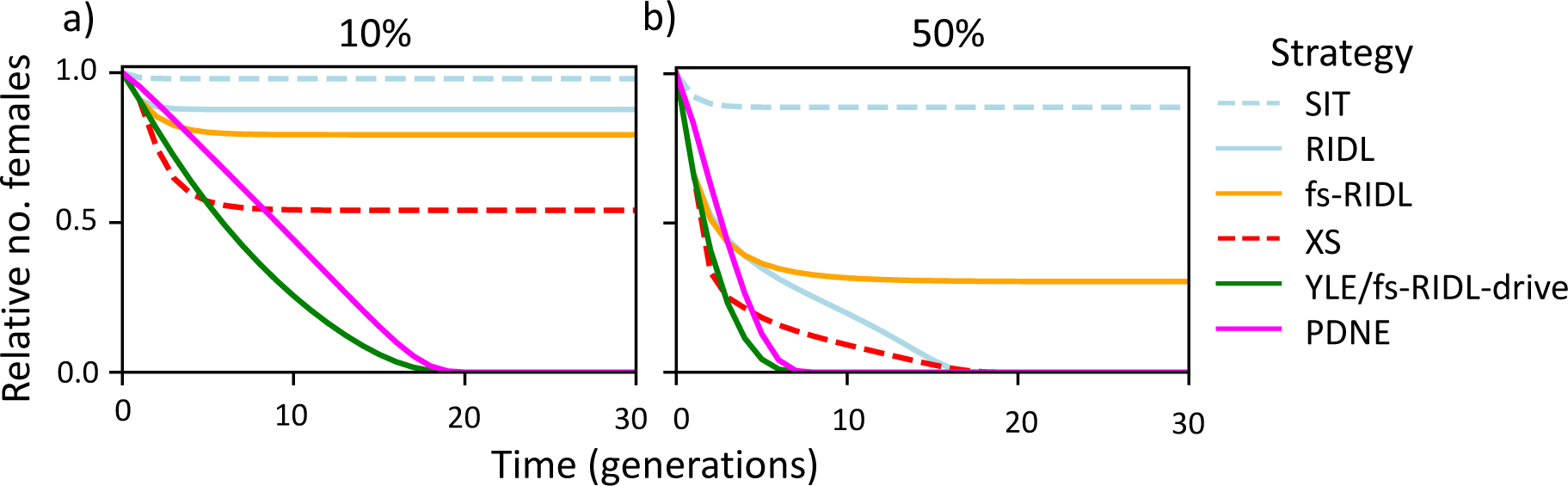
Time course simulations of the relative number of females following repeated releases each generation of males carrying different constructs at **(a)** 10% and **(b)** 50% of the original male population size. Strategies include release of: sterile males that result in death of offspring before density dependent mortality (SIT, blue, dashed), males homozygous for either a dominant lethal allele which causes death after density dependent mortality in both sexes (RIDL, blue, solid) or only females (fs-RIDL, orange, solid), a sex-ratio distorter (XS, red, dashed) or males carrying one copy of the Y-linked editor (green, solid) or PDNE (pink, solid) assuming all costs cause death after density dependent mortality. All strategies are modelled with idealised parameters in a population with an intrinsic rate of increase of 6. The PDNE was simulated using a single locus design, however in idealised conditions the results are equivalent for two-locus implementations.

**Fig. 3.**
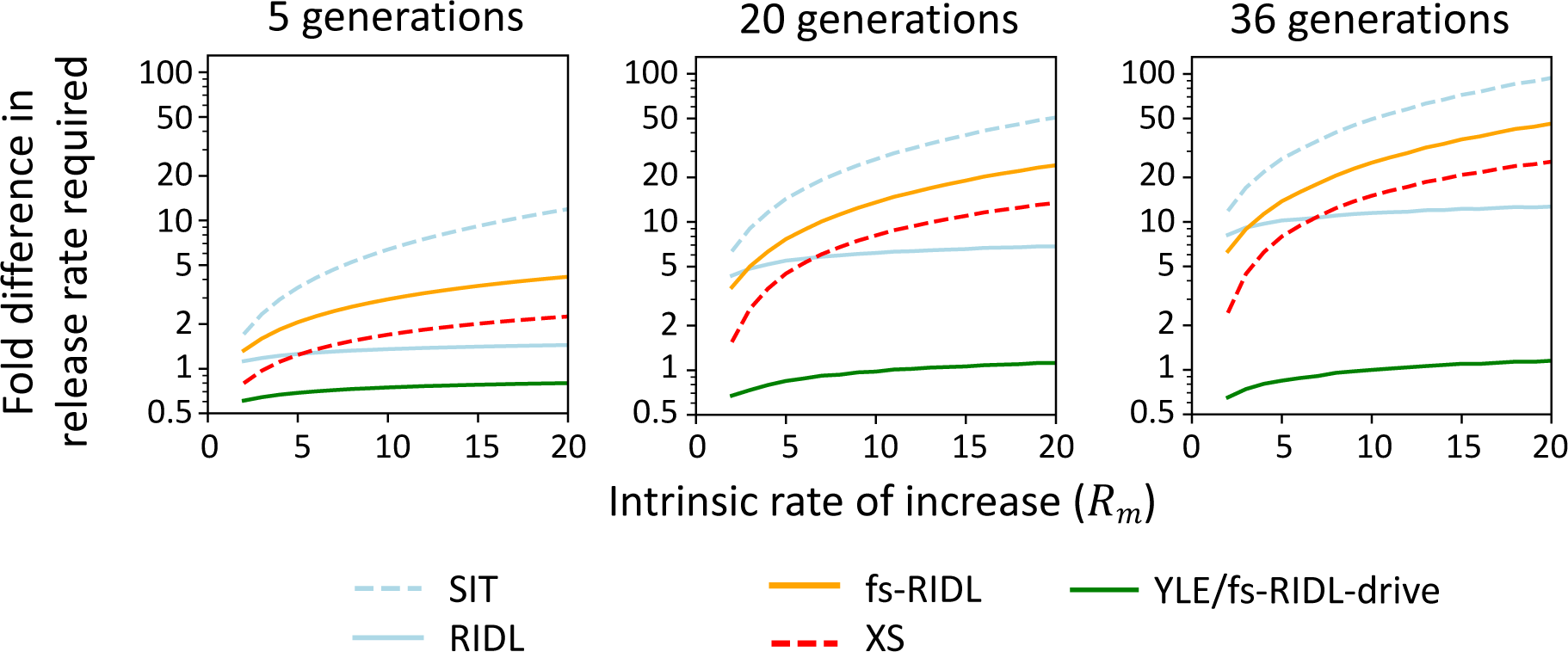
The fold-difference in the number of males required for release each generation to suppress a population with a range of intrinsic growth rates by 95% for a range of alternative strategies (described in Fig. 2) compared to the PDNE. From left to right plots show varying time frames in which the 95% suppression is required. Fold-differences are calculated by dividing release rates for each strategy by the release rate required for the PDNE to achieve equivalent levels of suppression. All strategies are modelled with idealised parameters.

**Table 1.** The number of released PDNE heterozygous males (as a proportion of the starting male population) required to suppress the level of females by 67%, 95% or 99% within 5 or 36 generations in a population with an intrinsic growth rate of 2, 6 or 20. Square brackets show the fold reduction in release rates compared to the sterile insect technique. All model parameters are idealised.

| | Intrinsic rate of increase ($R_m$) | | |
| --- | --- | --- | --- |
|  | 2 | 6 | 20 |
| Level of suppression within 36 gens |  |  |  |
| 67% | 0.017 [16] | 0.03 [43] | 0.036 [135] |
| 95% | 0.023 [12] | 0.042 [31] | 0.052 [94] |
| 99% | 0.025 [11] | 0.044 [30] | 0.056 [87] |
| Level of suppression within 5 gens |  |  |  |
| 67% | 0.217 [2] | 0.303 [6] | 0.344 [20] |
| 95% | 0.481 [2] | 0.685 [4] | 0.819 [12] |
| 99% | 0.693 [1] | 0.992 [3] | 1.228 [9] |

### 2.3. Robustness of performance and behaviour

In practice, it will not be possible to engineer a construct which behaves in the idealised way we have modelled and therefore it is important to explore the impact of possible imperfections on its performance and behaviour. The two most important considerations are, first, how various imperfections may affect the efficiency of the system – the release rates required – and, second, whether there are any imperfections that could cause the construct to drive and no longer remain localised. Considering first the effects on efficiency, Figure 4a-c shows the release rates required to suppress a population with an *R_m_* of 2, 6 or 20 by 95% within 36 generations whilst varying the editing parameters, keeping all other parameter values as ideal. When parameters associated with creation of the dominant edit are suboptimal, such as the editing rate and fitness costs of the edit, the overall strength of drive reduces, tipping the balance in favour of selection and reducing the constructs persistence. This, along with reduced load due to fewer or weaker dominant mutations being created increases the release rates required to achieve the desired level of suppression. Interestingly, similar suppression efficiencies can be achieved if the edits created only affect female fitness and in some cases this can even provide a gain in efficiency (Fig. 4 dashed lines). The PDNE strategy is most tolerant of suboptimal fitness costs of the edit in homozygotes (Fig. 4b), where release rates are little affected by up to 10% reduction in homozygote costs when suppressing a population with an Rm of 20, increasing to 50% in populations with an *R_m_* of 2, although with greater costs efficiency rapidly declines and the strategy fails. In *Drosophila* most recessive lethal alleles have at least some small heterozygous fitness effects (Crow and Temin, 1964) and in previously engineered gene drives leaky expression of Cas9 causing editing in the soma has also led to unintended fitness costs in heterozygotes (Hammond et al., 2016, Fuchs et al., 2021). If there is increased lethality in individuals heterozygous for the PDNE, either due to costs associated with disruption of the construct insertion site or expression of the editor, selection against the construct is increased and consequently release rate requirements are greater (Fig. 4d).

**Fig. 4.**
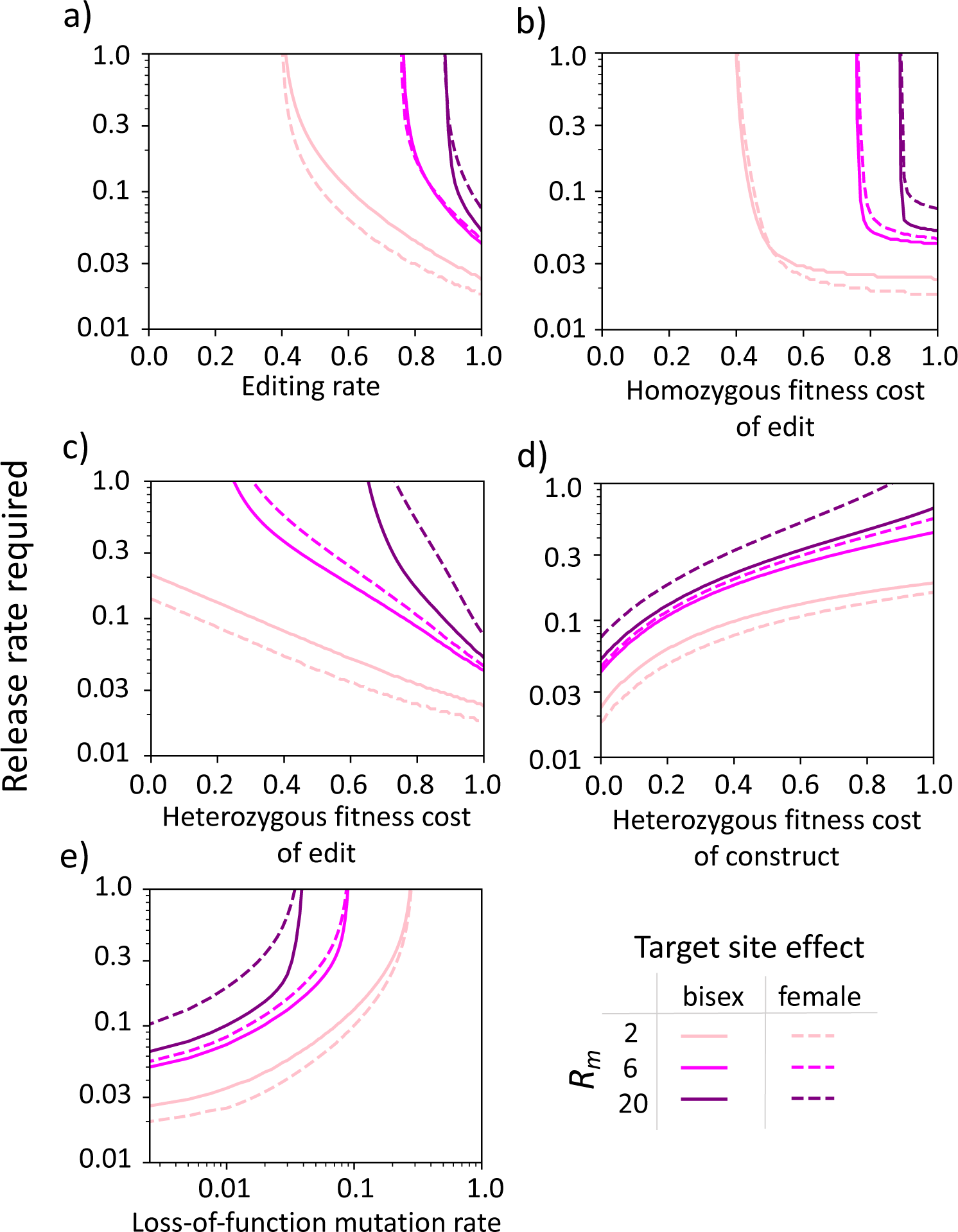
The release rates required to suppress the relative number of females in a population by 95% within 36 generations when varying parameters associated with the construct activity, fitness costs and stability, including **(a)** the probability the construct creates a dominant edit, **(b)** the homozygous fitness cost of the edit, **(c)** the heterozygous fitness cost of the edit, **(d)** the heterozygous fitness cost of the construct itself, and **(e)** the probability each construct component acquires a loss-of-function mutation each generation. Release rates are shown for populations with intrinsic rates of increase of 2 (peach), 6 (magenta) and 20 (purple) and for designs in which the dominant edit created by the PDNE affects both sexes (solid lines) or only females (dashed lines). All other parameters are idealised.

To assess the possibility that some imperfection in the construct might lead to spread through a population and control no longer being localised, we first return to our analytical model to explore the conditions under which the construct can drive. In the case where all edits produced by the editor are fully dominant and penetrant lethals (i.e. the edit responsible for the driving force has maximal effect), the construct is not expected to drive as long as 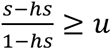 or *s* = 1, where *s* is the fitness cost in homozygotes, *hs* is the fitness cost in heterozygotes and *u* is the editing rate. Thus, as long as the recessive costs of the construct are also fully penetrant lethals (*s* = 1) drive can be prevented and the construct will not spread. Second, once released it is also possible for the components of the construct to acquire loss-of-function mutations that change the construct’s behaviour. To check whether any deletion-derivative constructs are able to spread we simulated releases of the construct with 1% chance of each molecular component (Cas9 and gRNA) losing function each generation. Our results show that none of the alleles containing non-functional components can drive from low frequency and all remain below the release frequency of the original PDNE after single or repeat releases (Supp. Fig. 2). Rather, loss-of-function mutations reduce the efficiency of the strategy since genomic editing is not possible without both the Cas9 and gRNA, leading to fewer dominant edits being created, reducing both persistence and load (Fig. 4e).

### 2.4. Boosting efficiency

Though our design offers significant improvement in efficiency over alternative strategies, further reductions in release rate requirements can be made by releasing the construct alongside one or more “booster” constructs that facilitate a temporary increase in the frequency of the first (DiCarlo et al., 2015, Esvelt et al., 2014). One approach is to release a booster which allows the effector construct to home in its presence but itself is inherited Mendelianly and is therefore lost over time rendering boosting temporary. A homing-based booster could be implemented by simply releasing a second construct containing a gRNA engineered to guide the Cas9 expressed from the PDNE to cleave the WT version of the PDNE insertion site, relying on the cell’s homology directed repair machinery to home the PDNE. Fig. 5b shows a time series simulation of a single release of individuals heterozygous for the PDNE and one copy of the homing-based booster at an unlinked locus. The PDNE (blue) increases in frequency in the presence of the booster (grey) which gradually disappears. With repeated releases, a homing-based booster can reduce the numbers of males needed to be released each generation by up to 10-fold (Supp. Fig. 3 and 4). For instance, releasing the PDNE alongside two copies of a booster reduces the required release rate to achieve 95% suppression of a population with an *R_m_* = 6 within 36 generations from 4.4% to 0.4%. Compared to an optimal version of SIT this approach can be three orders of magnitude more efficient (Supp. Fig. 3). Greater persistence of the PDNE means that boosting it, rather than alternative load-inducing constructs, can be considerably more impactful, offering up to a 100-fold increase in efficiency compared to boosting an X-shredder or fsRIDL construct (Supp. Fig. 4). In species where homing rates are low, an alternative approach could involve a cleave-rescue booster which increases the frequency of the effector by killing individuals which do not inherit it and therefore providing a fitness advantage (Fig. 5b, e).

**Fig. 5.**
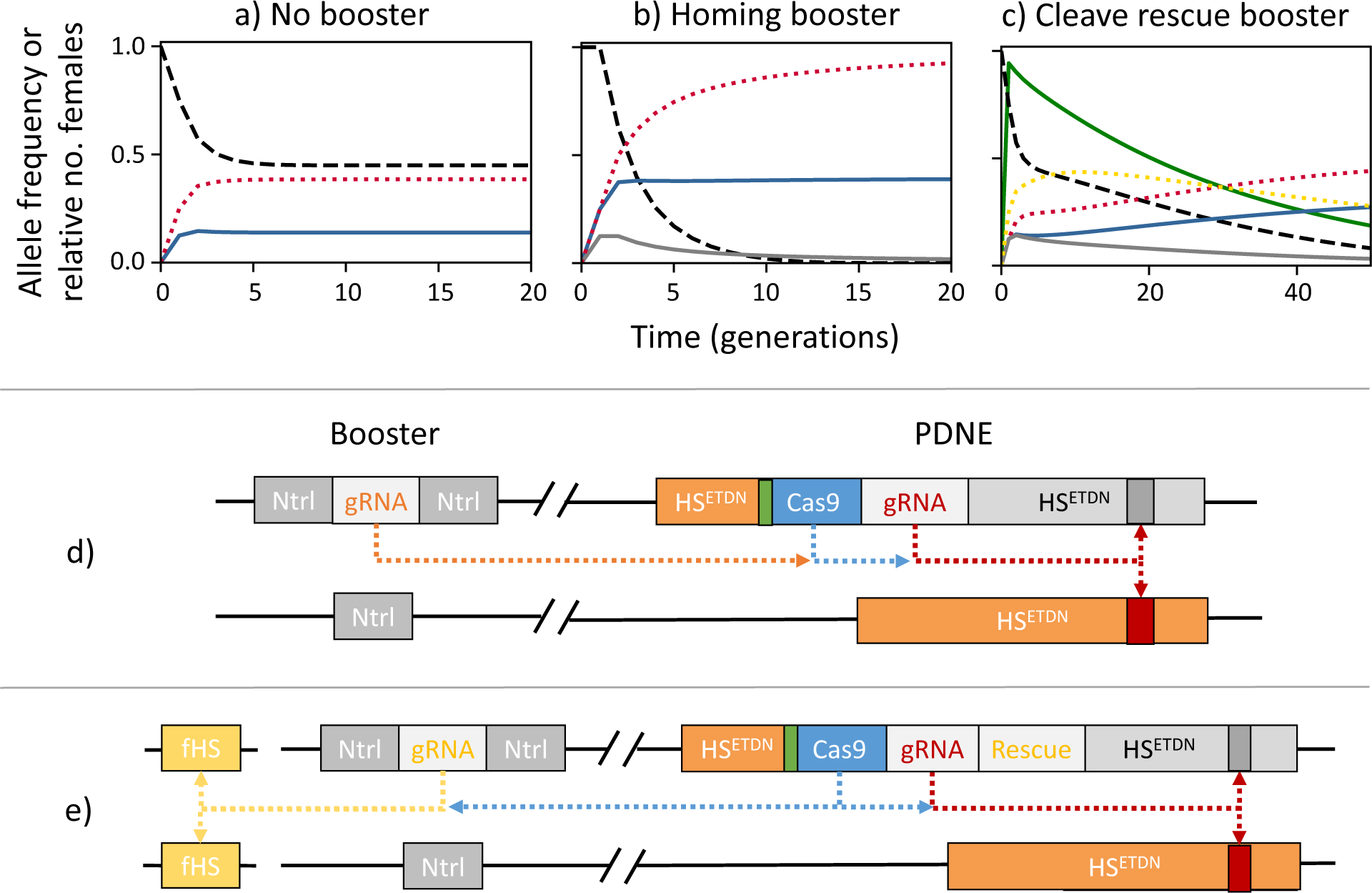
Time series simulations of single releases of males heterozygous for an idealized PDNE with or without homing or cleave-rescue booster constructs at 100% of the initial male population **(a-c)** and possible CRISPR-based molecular configurations **(d and e)**. When the PDNE is released in males that also carry an unlinked homing-based booster that allows the PDNE to home in its presence **(b)**, the PDNE (blue, solid) reaches a greater frequency than it would do without the booster **(a)**. Since the booster (grey, solid) is Mendelianly inherited, it is gradually lost over time and therefore boosting remains temporary. The higher frequency reached by the PDNE causes a greater load on the population and consequently causes a greater level of suppression (black, dashed). Panel **(c)** shows an alternative approach where a female-specific PDNE (blue, solid) is released in males which also carry a cleave-rescue booster (grey, solid) that creates edits in an unlinked gene to disrupt its function (yellow, dotted). In this example, the female-specific HS edits created by the booster act to reduce the fitness of individuals (or their descendants) who would have otherwise survived the PDNE-created female-specific HI mutation (i.e. males). If the PDNE is designed such that it rescues the function of these edits the presence of the booster will increase the selective advantage of the PDNE over the wild-type, increasing its frequency. Consequently, the PDNE reaches greater frequencies and inflicts a greater load on the population, increasing the level of suppression achieved. In this example, the two constructs are linked (r=0.05). High correlation between the two constructs (green, solid) facilitates persistence of the booster, increasing boosting. Over time recombination breaks down the correlation, and the booster decreases in frequency when inherited in individuals who do not carry protective PDNE and who succumb to the fitness effects of either of the costly edits. Diagrams **(d)** and **(e)** show possible molecular configurations of the two boosting strategies. A homing-based booster can be engineered using a gRNA targeting the insertion site of the PDNE and can be linked or unlinked to the PDNE construct. A cleave-rescue booster can similarly be constructed by 1) engineering the booster to contain a gRNA which guides the Cas9 expressed from the PDNE to create a costly edit at a second functional target gene and 2) modifying the PDNE to contain a cleavage resistant recoded copy of the target gene that rescues function. To facilitate comparison, all simulations involve a PDNE that induces female-specific dominant edits, although the homing-based booster can also effectively boost a bi-sex design too.

## 3. Discussion

Given that the inefficiency of SIT is due to the disappearance of the effecter allele within a single generation, an obvious way to improve efficiency is to increase its persistence. One previous proposal in this direction is fs-RIDL, in which the lethal effects of the genetic modification released is limited to females and therefor can persist across generations through males (Thomas et al., 2000). In this paper we have proposed a way of further increasing persistence of a costly allele by counterbalancing negative selection with genetic drive such that the construct is selectively neutral despite imposing a reproductive load. Once released into a population, the combination of load and selective neutrality allows our construct to act, at least in the idealised case and in the absence of immigration, like a genetic perpetual motion machine, suppressing the population indefinitely without the need for additional releases. This behaviour is expected to result in geographically confined population suppression with considerably lower release requirements than the currently utilised SIT and a range of alternative candidate strategies.

Our proposed design relies on three key features involving an autosomal construct that (1) causes recessive lethality in both sexes, (2) encodes a genome editor creating female-specific or bi-sex dominant lethal edits, and (3) provides some protection against the action or consequences of the editor. Although in principle the design could be implemented in many ways, it is possible to engineer this genetic strategy using current molecular tools, and we have proposed simple one- and two-locus CRISPR-based cleave-rescue approaches. Other strategies with a similar molecular configuration but which differ in their dynamics include cleave-rescue (ClvR) or toxin-antidote recessive embryo (TARE) designs for population modification (Oberhofer et al., 2019, Champer et al., 2020b) and strategies for suppression that exhibit drive (TADE) (Champer et al., 2020a). Other strategies with non-driving, selectively neutral dynamics similar to our design but which take a different molecular approach include YLEs and fs-RIDL-drives (Burt and Deredec, 2018). YLEs are genomic editors which generate costly edits that affect females and achieve selective neutrality due to being located on the Y chromosome where they are hidden from selection in females. In contrast, the fs-RIDL-drive construct is autosomal and causes dominant costs in females that inherit it. Selective neutrality is achieved similarly to the PDNE, balancing selection against the dominant lethality in females with homing-based genetic drive in males. Although both strategies offer comparable efficiency to the PDNE, the YLE requires integration into, and germline expression from, the Y-chromosome, which can be difficult to achieve (Turner, 2007, Bernardini et al., 2014, Taxiarchi et al., 2019) and also restricts the approach to species with a Y-chromosome. The efficiency of the fs-RIDL-drive strategy depends on high homing rates which in some species are difficult to achieve (Grunwald et al., 2019, Pfitzner et al., 2020, Weitzel et al., 2021). Both strategies also require genes in which female-specific dominant mutations can be made, which are rare and can be difficult to work with, limiting target site choice. Our proposed design evades these difficulties since it does not require homing, can be built using autosomal loci in species with or without sex-chromosomes and does not require sex-specific fitness effects, and therefore may be easier to engineer.

Our single-locus PDNE configuration (Fig. 1c lower) offers a particularly attractive option for construction since it prevents the need to include a recoded copy of a gene within the construct, which in some cases has been difficult to achieve (Chen et al., 2023a). The design relies on integrating the construct into a haplo-sufficient gene needed in both sexes in which it is possible for the genomic editor to create a dominant negative mutation. The same approach would also work with haplo-sufficient genes which are editable to create dominant gain of function mutations, in which the protein takes on a new function that causes sterility or lethality. A search of FlyBase indicates there are 45 genes in *Drosophila melanogaster* with both dominant lethal and recessive lethal mutations, providing a starting point to identify a suitable gene. Although the most efficient target gene would be one which affects survival after density-dependent mortality occurs, genes which affect survival early in development or impact fertility can also offer effective suppression (Supp. Fig. 5). *doublesex* is a noteworthy gene involved in insect sex-determination for which both bi-sex recessive and female-specific dominant sterile mutations have been identified (Chen et al., 2023b, Tolosana et al., 2024), and also forms the basis of a range of emerging synthetic genetic control strategies for suppression (Kyrou et al., 2018, Yadav et al., 2023, Simoni et al., 2020, Kandul et al., 2019). Although in our design we propose inserting the construct upstream of the target site to prevent expression of the dominant negative or gain of function edit, as long as the haplotype which contains the editor also contains a disrupted version of the haplo-sufficient gene, the construct could alternatively be located near to, but downstream of the edit, inside or outside of, or within a nearby haplo-sufficient gene. In cases where its insertion does not prevent expression of the edited transcript, the editor target site located in the same haplotype could simply be deleted, preventing the edit from being created and thus providing an alternative means of protection without incorporating a recoded gene into the construct.

In our alternative two-locus molecular configurations, the construct is inserted into a haplo-sufficient gene and the editor engineered to target a separate gene elsewhere in the genome (Fig. 1c upper). This approach would allow for the editor to create a dominant edit by knocking out an HI gene, therefore broadening the potential choice of target sites. 818 genes in FlyBase are annotated as having an amorphic or loss of function allele giving a recessive lethal phenotype, suggesting ample choice for insertion sites. Cook et al. (2012: Table 2) list 43 genes in *Drosophila melanogaster* that are haplolethal or haplosterile, plus another six regions of the genome that appear to have a haplolethal or haplosterile within them but the exact gene had not yet been identified, offering a range of putative target genes. If the two genes are linked, protecting the construct against the edit could be achieved by recoding the target gene in situ, whereas if they are unlinked, it would need to form part of the construct.

Regardless of the precise configuration, effort should be made to prevent the production of unintended functional or recessive edits by the editor, since these are likely to hinder the efficiency of the strategy by reducing the strength of drive. Following from previous suggestions, the probability of creating unwanted edits could be mitigated by targeting a highly conserved gene or sequence, or by using multiple gRNAs (Kyrou et al., 2018, Oberhofer et al., 2019, Yang et al., 2022, Champer et al., 2020c). In addition, one could increase the likelihood of making a precise sequence change by using prime- or base-editing (Marr and Potter, 2021, Bosch et al., 2021) or by releasing the desired sequence alongside the construct for use as homology directed repair template. Next, to guarantee the construct and its impacts remain localised it would be good to ensure that the recessive costs of the construct are both evolutionarily stable and 100% penetrant. Our proposed designs induce recessive lethality or sterility by inserting the construct into or linking it to a disrupted allele of a HS gene required for survival or fertility. With this design loss-of-function mutations in any of the construct components will not affect the strength of selection, and therefore none of the construct derivatives are expected to drive. An alternative molecular design could involve inserting the construct into a neutral site and including a second gRNA which creates recessive edits at a separate HS gene, however loss-of-function mutations in this gRNA would decrease selection and therefore create deletion derivatives more likely to spread. Furthermore, to prevent drive it will be important to ensure that the editor does not home when the edit is created, whether by ensuring sufficient distance between the editor and target site, or by using prime- or base-editing. Some studies have found that parental deposition of the Cas9 and gRNA can either hinder or enhance drive (Hammond et al., 2021a, Champer et al., 2020b, Oberhofer et al., 2019). In our designs we expect deposition to increase selection, rather than drive, since cleavage of the WT target allele in construct-bearing offspring would increase selection against the construct, whereas cleavage in non-construct-bearing offspring would be redundant since they already inherit an intended dominant lethal mutation. Lastly, wild populations may deviate from the random mating assumption made so far, instead exhibiting some degree of inbreeding. In these cases, we would not expect drive to occur since selection against the recessive costs of construct would be expected to increase (due to homozygotes for the construct being more often than in Hardy-Weinberg frequencies) and therefore the construct is more likely to decline in frequency.

Previous studies have demonstrated that for strategies involving two constructs at separate loci the degree to which the two loci are linked can influence their dynamics, and for this reason genetic distance can be tuned to control persistence and efficiency (Willis and Burt, 2021, Oberhofer et al., 2021b). This also applies to our designs involving releasing the PDNE alongside a homing or cleave-rescue booster, though the quantitative details between designs differ. Interestingly, we find that a homing booster can be located within the same gene as the PDNE, essentially forming part of the same construct, and boosting will remain temporary as long as the booster does not cause itself to home. This design may be particularly attractive from a regulatory perspective, since it involves only a single construct, or for geographically restricted gene drive trials since it exhibits drive temporarily. In theory, our boosted designs could also form the basis of a low-threshold geographically restricted double drive, and due to their selective neutrality may offer increased resilience to resistant alleles compared to alternative designs (Willis and Burt, 2021, Geci et al., 2022). In principle further control of our construct’s dynamics could be achieved by using seasonally important genes (Oberhofer et al., 2021a) or environmental application of chemicals that affect the fitness effects of the construct. If the gene used to induce recessive costs is only needed periodically or not needed in presence of chemical, then during specific times or in the presence of the chemical drive will outweigh selection and allele frequencies will temporarily increase. Alternatively, if such a gene was used to induce the dominant costs, selection would outweigh drive, and the construct would be purged from population under certain conditions. Finally, choosing genes with functions that can be reconstituted in the laboratory might facilitate large scale production by allowing for pure breeding lines and alleviating the need for time-consuming screening.

In summary, our proposed approach expands the current genetic biocontrol toolbox offering novel cleave-rescue designs for achieving selectively neutral localised suppression. While most current synthetic homing and toxin-antidote constructs mimic naturally evolved drive systems, our design is not expected to have evolved naturally, and rather is a product of a novel combination of available molecular building blocks motivated by end use. Given the success in using CRISPR/Cas9 to create alternative constructs it is conceivable that the construct(s) can be built using currently available molecular tools and may be easier to construct than current alternatives. Which of our proposed molecular configurations would be most applicable to different species and control goals will depend on the biology of the target population, the way in which harm is inflicted by the pest and the genes and technologies which are available within the species. In future, it will also be useful to further compare the resilience of each design to different types of resistance, unintended fitness costs and loss-of-function mutations, and to develop more context- and species-specific models to explore optimal rearing and release strategies and compare the potential for dispersal from the release site across designs.

## Acknowledgements

We would like to thank Federica Bernardini and Ignacio Tolosana for useful discussions. This work was supported by the Bill & Melinda Gates Foundation and Open Philanthropy

