## Supplementary Material for "Engineering drive-selection balance for localised population suppression with neutral dynamics"

### 1. Supplementary figures

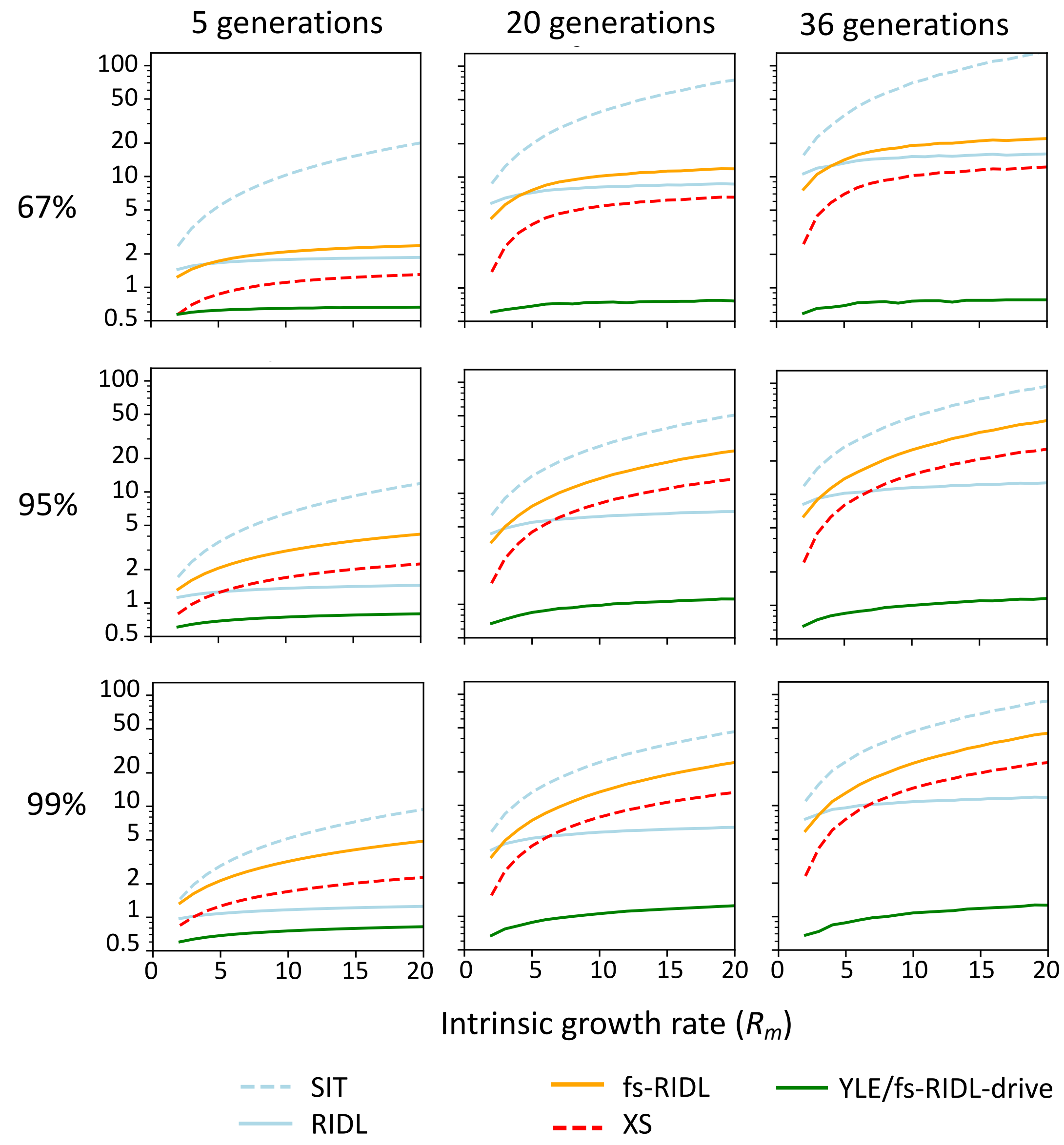


**Supp. Fig. 1** – The fold-difference in male release rates required to suppress a population by 67%, 95% and 99% using a range of strategies (as in Fig. 2) compared to a PDNE as a function of the intrinsic growth rate of the target population. From left to right plots show varying time frames within which the level of suppression is achieved. Parity to the PDNE is indicated by the grey dotted line. All strategies are modelled with idealised parameters.

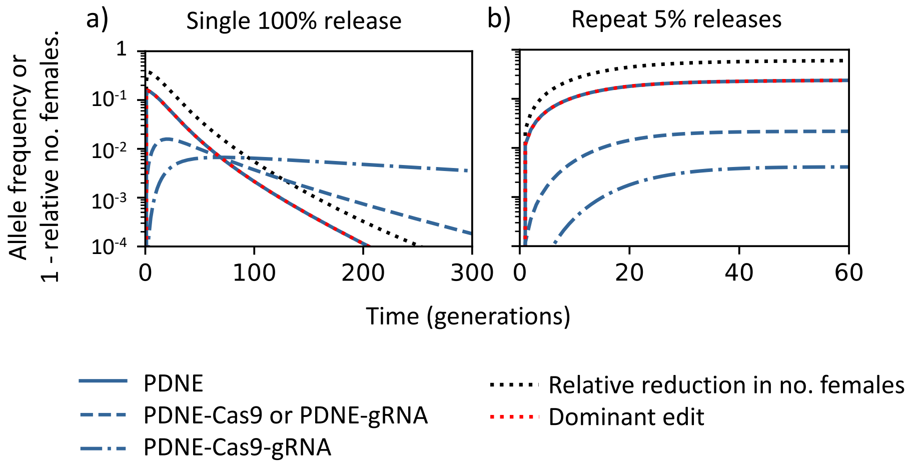


**Supp. Fig. 2** – Time series simulations of **(a)** single and **(b)** repeated releases of males heterozygous for an idealized PDNE where loss-of-function mutations occur in each construct component (Cas9 and gRNA) at 1% per generation. After a single release in the first generation the intact construct (PDNE, blue solid) creates edits (dominant edit, red dotted) in the first generation after release causing an increase in the relative reduction in females (black, dotted). In subsequent generations the intact construct decreases, owing to the construct accumulating loss-of-function mutations in either the Cas9 or gRNA (blue dashed) or both (blue dot-dashed). None of the derivative constructs generated through loss-of-function mutations are able to drive and each remain below the frequency of the released construct in both single and repeat release regimes.

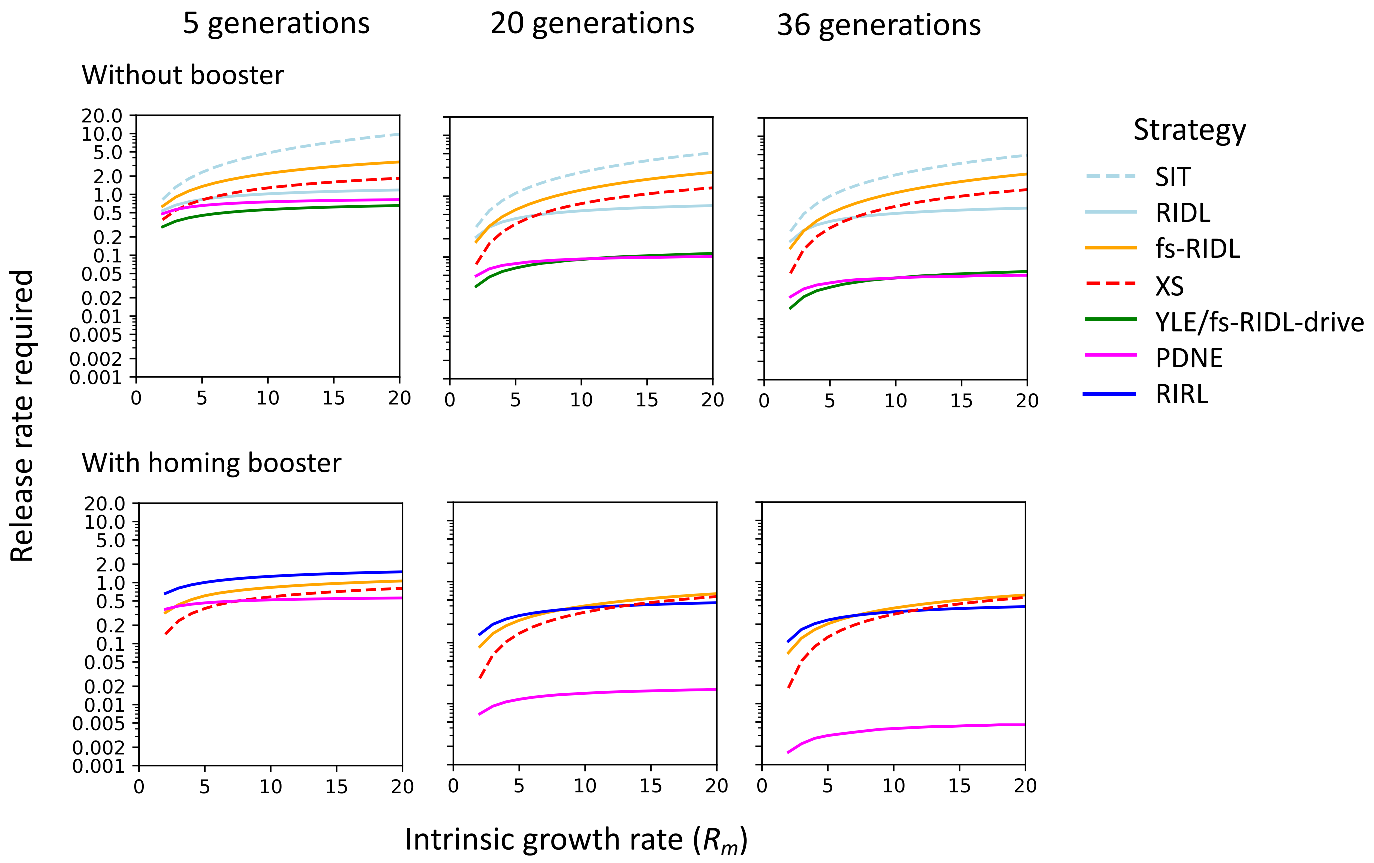


**Supp. Fig. 3** – Release rate requirements to suppress a population with a range of intrinsic growth rates by 95% using a range of suppression constructs (as in Fig. 2) released with (upper) and without (lower) a booster. From left to right plots show varying time frames in which the 95% suppression is achieved. For all scenarios we assume release of two copies of the booster unlinked to the construct. Note that since no offspring who inherit the SIT or RIDL constructs survive, the presence of a booster makes no difference to release rate requirements (not shown). Additionally, since the YLE is on the Y-chromosome and the fs-RIDL-drive homes itself, neither can be boosted via homing (not shown). Boosting in individuals carrying a recessive lethal (RIRL) was also modelled (b, dark blue solid), equivalent to the PDNE without the driving force, which, in the absence of the booster, is unable to suppress the population by 95% within reasonable release rates (not shown).

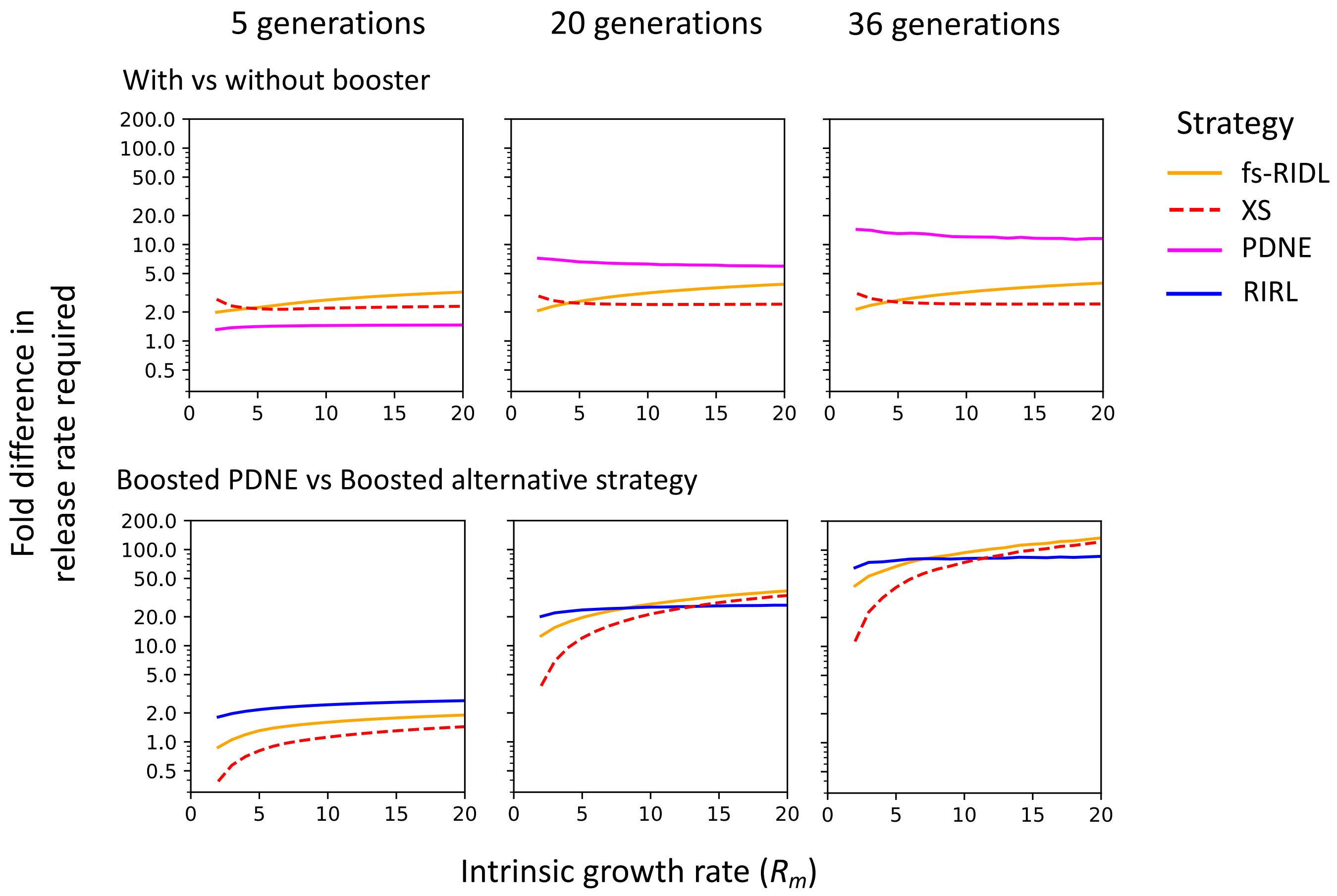


**Supp. Fig. 4** – The fold-difference in male release rates required for strategies (as in Supp Fig. 3) when comparing each strategy with and without a homing booster (upper) or comparing different boosted designs to the boosted PDNE (lower). All strategies are modelled with idealised parameters. Released males are homozygous for the booster which is unlinked from the PDNE.

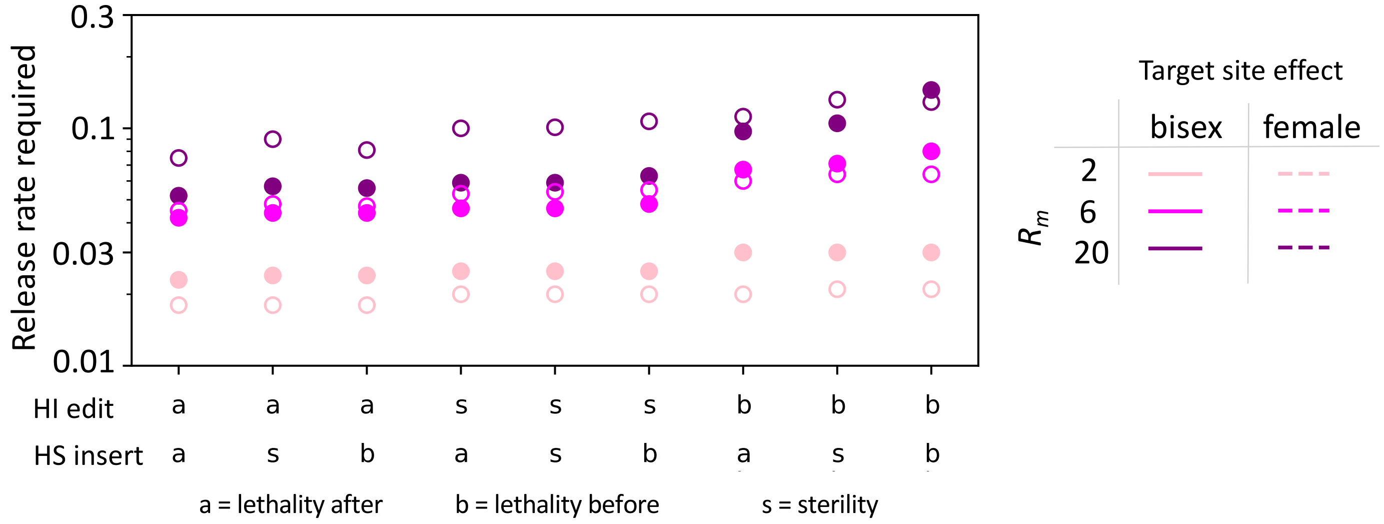


**Supp. Fig. 5** – The release rates required to suppress the number of females in a population by 95% within 36 generations when varying whether the dominant edit and recessive disruption causes sterility (s) or lethality before (b) or after (a) density dependent mortality. Release rates are shown for populations with intrinsic rates of increase of 2 (peach), 6 (magenta) and 20 (purple) and for designs in which the dominant edit created by the PDNE affects both sexes (filled circles) or only females (open circles). All other parameters are idealised.

### 2. Considerations for two-locus implementations

Whether our proposed design is implemented using one or two loci, the impact of suboptimal parameters associated with the edit or unintended costs in heterozygotes on the efficiency of the strategy is unchanged, at least when all other parameters are idealised. However, when constructing designs involving two loci it is important to consider the effect of linkage between the loci. For designs where the editor creates haplo-insufficient edits in a different gene to which it is inserted, and a recoded copy of that gene is located within the construct (Fig. 1c, upper), linkage between the loci can reduce efficiency when editing rates are suboptimal (Supp. Fig. 6). In contrast, in designs where the editor targets a gene closely linked to its insertion site and the recoded copy of the HI gene is in its native genomic location, tight linkage between the construct and the HI gene is vital to maintain impact (Supp. Fig. 7). For constructs that contain a recoded copy of the construct (whether in the construct or linked to it) it is possible that rescue functionality is suboptimal or that the rescue element acquires loss-of-function mutations that prevent rescue. Regardless of linkage, suboptimal rescue is expected to reduce the strength of drive since the construct will be subject to (at least some) selection against the dominant edit. This would not result in drive of the construct and instead decrease efficiency of the design, although quantifying the magnitude of this effect would require further analysis that is outside the scope of this work.

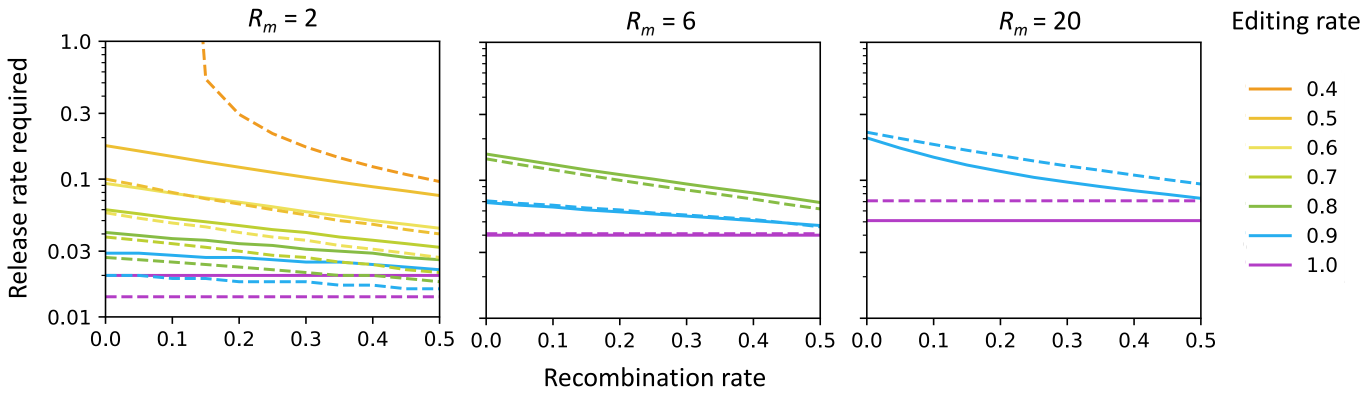


**Supp. Fig. 6** – The release rates required of males carrying a PDNE implemented using two loci to suppress the level of females in a population by 95% of its original size within 36 generations as a function of the recombination rate between the PDNE and its target site and the mutation rate for populations with an intrinsic growth rate 2, 6 or 20. The fitness effects of the edit created by the PDNE affect either both sexes (solid lines) or only females (dashed lines) and are assumed to cause lethality after density dependent mortality. Recall from figure 4 that for some editing rates suppression to the desired level is not achievable with release rates less than one, therefore not all editing rates are relevant to all $R_m$ values used. All other parameters are assumed to be idealised.

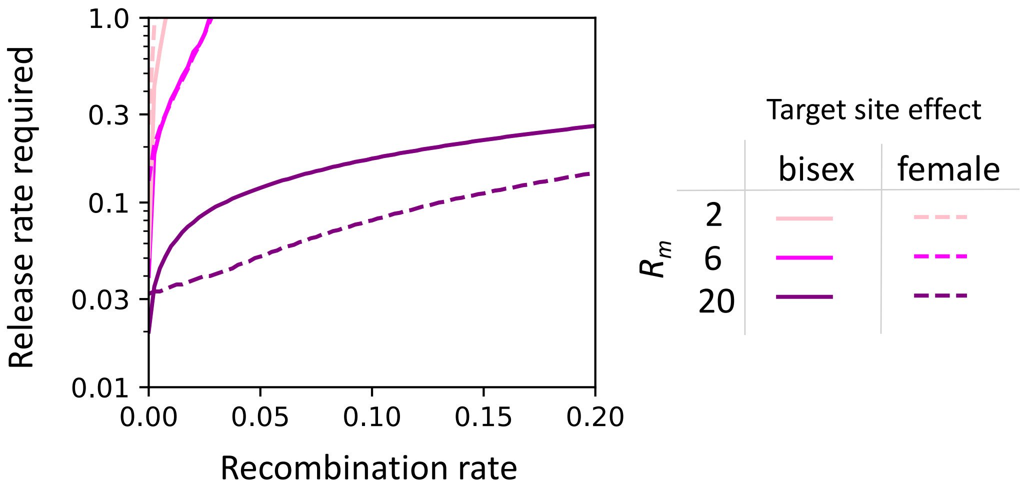


**Supp. Fig. 7** – The release rates required to suppress the number of females in a population by 95% within 36 generations using a 2-locus implementation of our design where the construct causes recessive fitness costs and creates knockout mutations in a linked haplo-insufficient gene. Rescue is achieved by releasing the construct linked to a fully functional version of the haplo-insufficient gene that is resistant to cleavage.
